# Interferon Regulatory Factor 5 (IRF5) Interacts with the Translation Initiation Complex and Promotes mRNA Translation During the Integrated Stress Response to Amino Acid Deprivation

**DOI:** 10.1101/163998

**Authors:** James D. Warner, Mandi Wiley, Ying-Y Wu, Feng Wen, Michael Kinter, Akiko Yanagiya, Kandice L. Tessneer, Patrick M. Gaffney

## Abstract

Interferon Regulatory Factor 5 (IRF5) plays an important role in limiting pathogenic infection and tumor development. Host protection by IRF5 can occur through a variety of mechanisms including production of type I interferon and cytokines as well as the regulation of cell survival, growth, proliferation, and differentiation. While modulation of these cellular processes is attributed to IRF5 transcription factor function in the nucleus, emerging evidence suggests that IRF5 may also retain non-transcriptional regulatory properties within the cytoplasmic compartment. Consistent with this notion, we report the ability of IRF5 to control gene expression at the level of mRNA translation. Our findings demonstrate that IRF5 interacts with the translation initiation complex in the absence of the m^7^GTP cap-binding protein, eIF4E. We observed that under nutrient deprivation-induced cell stress, IRF5 promoted mRNA translation of the master integrated stress response (ISR) regulator, Activating Transcription Factor 4 (ATF4). Enhanced ATF4 protein expression correlated with increased levels of downstream target genes including CHOP and GADD34 and was associated with amplification of eIF2α de-phosphorylation and translational de-repression under stress. The novel mechanism we describe broadens our understanding of how IRF5 regulates gene expression and may govern diverse cellular processes in the absence of stimuli that trigger IRF5 nuclear translocation.

## INTRODUCTION

Protein synthesis is a process essential for cell survival, however it is a function that renders the cell vulnerable to changes in nutrient availability and viral replication. As a result, safeguards are in place to ensure coordination between cell activity and environmental conditions. Many of these protective measures surround the rate-limiting, initiation step of mRNA translation.

During mRNA translation initiation, the ribosomal subunits are recruited in stages to the translation start site by eukaryotic initiation factors (eIFs) that form the translation initiation complex (TIC). Central to this process is the m^7^GTP-binding protein, eIF4E, which binds the 5’-cap of mRNA transcripts; eIF4A, a helicase that unwinds RNA secondary structure; and eIF4G, a scaffolding protein that binds both eIF4E and eIF4A as well as other factors important for mRNA translational regulation including the eIF3 complex (1-4). eIF3 is responsible for TIC recruitment of the 43S preinitiation complex which is comprised of the 40S ribosomal subunit, eIF2α bound by GTP, and the initiator methionyl-tRNA (Met-tRNAi) (5,6). Upon recruitment, 40S ribosomal scanning commences and subsequent recognition of the start codon by Met-tRNAi results in eIF2-GTP hydrolysis and its release of Met-tRNAi as well as departure from the TIC along with eIF3 thereby facilitating 60S ribosomal recruitment and downstream protein synthesis (5,7).

eIF2-GDP can be recycled for additional rounds of translation initiation, but only following the GDP-for-GTP exchange catalyzed by the guanine nucleotide exchange factor, eIF2B (8). Additionally, four different mammalian eIF2α kinases (eIF2AKs) exist which can phosphorylate eIF2α on Ser51 and prevent guanine nucleotide exchange resulting in inhibition of translation initiation (9,10). These eIF2AKs include General Control Non-derepressible 2 (GCN2), Protein Kinase R (PKR), PKR-like Endoplasmic Reticulum Kinase (PERK), and Heme-Regulated Inhibitor (HRI) and each of these kinases act as an environmental sensor to different sets of stimuli (9,10). GCN2 is activated in response to unbound tRNAs and certain RNA viruses and can therefore provide protection against amino acid deprivation and infection, respectively, by conserving the supply of amino acids and shutting down the translational machinery required for viral replication (11-13).

In addition to inhibiting the mRNA translation rate, GCN2 (like other eIF2AKs) upregulates an adaptive gene expression program known as the integrated stress response (ISR) (14). Central to the regulation of the ISR is the transcription factor, ATF4. ATF4 promotes the transcription of many target genes including those involved in nutrient uptake and metabolism, protein folding, and cell survival as well as other transcription factors that further regulate a broad array of biological processes (15-19). ATF4 also directly promotes upregulation of GADD34, an eIF2α phosphatase that stimulates translational de-repression during cell stress and thereby facilitates protein synthesis of ISR-related genes (20).

Protein expression of ATF4 is tightly regulated and mRNA translation is paradoxically triggered by eIF2α phosphorylation (21). An upstream open reading frame (uORF) present within the ATF4 mRNA transcript hinders translation initiation at the ATF4 translation start site when levels of Met-tRNAi-eIF2-GTP (collectively known as ternary complex) are abundant (21). However, when less ternary complex is available, prolonged ribosomal scanning occurs allowing for uORF bypass and translation of the ATF4 coding sequence (21).

IRF5 is a transcription factor that responds to pathogenic infection, DNA damage, and death receptor (DR) signaling, and therefore may also be viewed as a regulator of certain forms of cell stress (22-26). Ligand binding by the pattern recognition receptors, Toll-like receptor (TLR) 7 and TLR9, as well as binding of DR4/5 by TNF-related Apoptosis-inducing Ligand (TRAIL) leads to downstream signaling which activates IRF5 (24,27,28). IRF5 activation results in dimerization and nuclear translocation where it promotes transcription of type I interferon (IFN), pro-inflammatory cytokine, and apoptosis-related genes (29,30). While the role of IRF5 in the nucleus is well characterized, a recent report described a cytoplasmic function of IRF5 unrelated to transcription thereby suggesting that more insight is needed into the way IRF5 regulates various cellular processes (31).

In support of this idea, we report here a novel interaction between IRF5 and multiple TIC components including eIF4G and eIF4A, but not the m7GTP binding protein, eIF4E. Furthermore, we demonstrate that IRF5 promotes mRNA translation of ATF4 during amino acid deprivation resulting in enhanced expression of ATF4 target genes, protein synthesis, and altered ISR gene signature. Our work provides a broader understanding of the way in which IRF5 influences cell stress responses.

## RESULTS

### IRF5 interacts with the translation initiation complex independently of eIF4E

Mass spectrometry analyses of IRF5 immunoprecipitation reactions suggested associations between IRF5 and several RNA-binding proteins implicated in mRNA translation including 40S ribosomal proteins, translation elongation factors, putative IRES-transacting factors (ITAFs), and RNA helicases **(Table S1)** (32,33). Amongst the RNA helicases identified was the translation initiation factor (TIF), eIF4A. Given the importance of translation initiation in the regulation of protein synthesis, we initially focused on identifying and characterizing interactions between endogenous IRF5 and members of the TIC (34).

We used cell lysates from multiple cell lineages to perform immunoprecipitation (IP) of endogenous eIF4G, eIF4A, and PABP to assess interaction with endogenous IRF5. Chemical crosslinking with 1% formaldehyde prior to cell lysis was utilized to prevent the formation of spurious interactions between IRF5 and TIFs *in vitro*. We observed that IRF5 co-immunoprecipitated with eIF4G, eIF4A, and PABP across all cell lines tested (**Fig. 1a-c**). We further confirmed these interactions by performing co-IP experiments using lysates (without crosslinking) derived from HEK293T cells co-expressing either HA-tagged eIF4G and FLAG-tagged IRF5 (full-length; IRF5-V5) **(Fig. 1d)**, or HA-tagged IRF5-V5 and either FLAG-tagged eIF4A **(Fig. 1e)** or FLAG-tagged PABP **(Fig. 1f)**. Interestingly, we were unable to detect any such interaction between IRF5 and the m^7^GTP-cap binding protein, eIF4E, despite robust pull-down of the target protein (**Fig. 1g-i**), the use of antibodies recognizing distinct eIF4E epitopes (**Fig. S1a**), or the use of m^7^GTP-coupled sepharose beads (**Fig. S1b**). Collectively, these data suggest that IRF5 interacts with the TIC independently of eIF4E.

**FIGURE 1.**
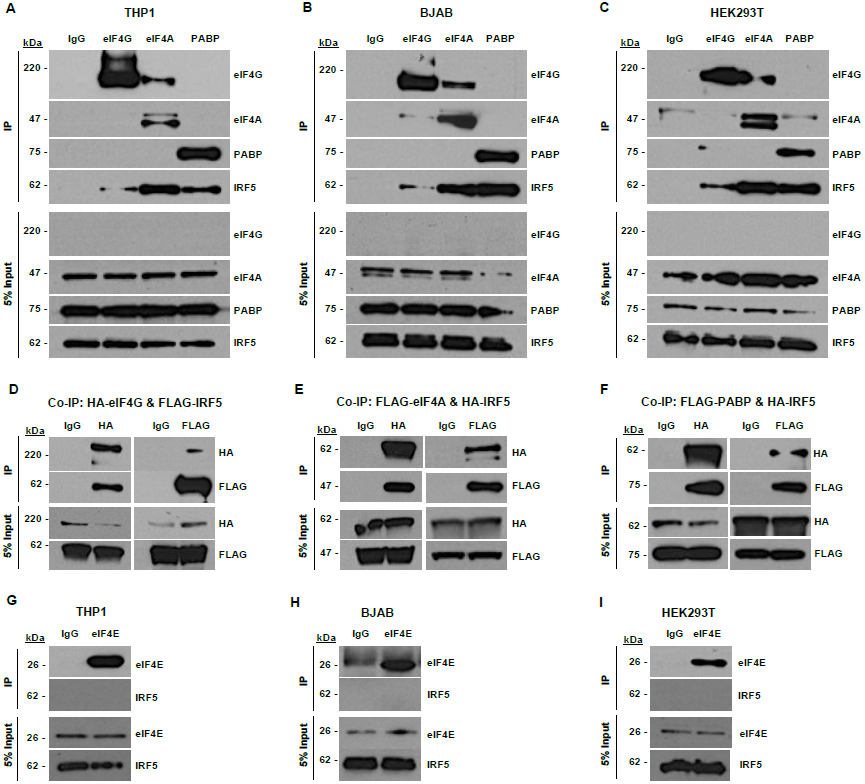
IRF5 interacts with the translation initiation complex. **A-C**, Endogenous eIF4G, eIF4A or PABP were immunoprecipitated from THP1 (A), BJAB (B), and HEK293T (C) cell lysates previously fixed with 1% formaldehyde and analyzed by immunoblotting as indicated. eIF4G levels were below the limit of detection within input samples due to insufficient protein abundance relative to percentage of input saved per reaction. **D-F**, HEK293T cells were co-transfected with the indicated plasmids, subjected to reciprocal co-immunoprecipitations, and analyzed by immunoblotting as indicated. **G-I**, Endogenous eIF4E was immunoprecipitated from THP1 (G), BJAB (H), and HEK293T (I) cell lysates previously fixed with 1% formaldehyde and then analyzed by immunoblotting as indicated. (**A-I**) All results are representative of n≥3 independent experiments.

IRF5 mRNA transcripts are expressed as different isoforms, but structural variability of IRF5 protein resides solely within exon 6 amongst the most common isoforms (35,36). Structural variability in exon 6 is based upon the combination of a 30-base pair insertion/deletion (indel) polymorphism and an alternative splice site that results in loss of the first 48 base pairs of exon 6 (**Fig. S2a**) (35). To investigate whether structural variability in IRF5 would impact IRF5 association with the TIC, we co-expressed HA-tagged eIF4G with four of the most common FLAG-tagged IRF5 variants (V1, V2/V6, V3/V4, and V5) in HEK293T cells (**Fig. S2b**). The scaffolding protein, eIF4G, which interacts with both eIF4A and PABP during translation initiation was used as a proxy during the examination of TIC-IRF5 variant interaction (37,38). We found no evidence that the IRF5 exon 6 indel or splice pattern influenced association with eIF4G **(Fig. S2b).** This may suggest that the interaction between IRF5 and eIF4G occurs outside of IRF5 exon 6 or is not sufficiently altered to disrupt such interactions.

### IRF5 binds the MIF4G domain of eIF4G

The eIF4G domains in which eIF4A, PABP, eIF3, and MAPK-interacting kinase 1 (MNK1) interact with the scaffolding protein are well characterized (39-43). To map the domain required for IRF5 and eIF4G interaction, we co-expressed plasmids encoding full-length or truncated HA-tagged eIF4G mutants **(Fig. 2a)** with FLAG-tagged IRF5-V5 in HEK293T cells. Co-immunoprecipitation data demonstrate that only eIF4G truncation mutants which contain the middle domain of eIF4G (MIF4G) possess the ability to bind IRF5 **(Fig. 2b)**.

**FIGURE 2.**
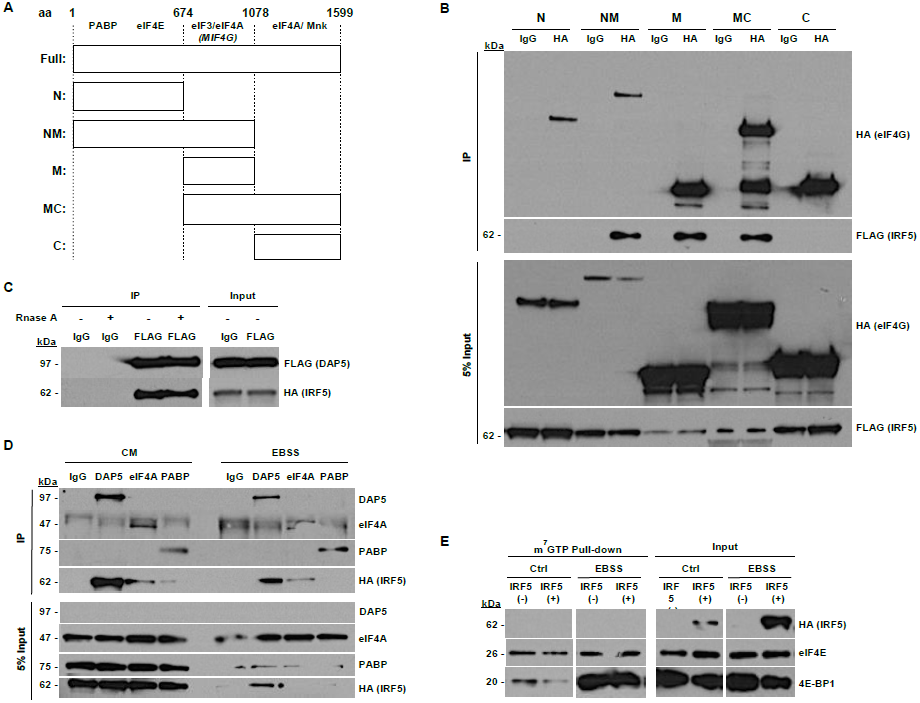
IRF5 interacts with the MIF4G domain of eIF4G. **A**, Diagram of HA-eIF4G constructs and known binding sites of interacting translation initiation complex partners. **B,** HEK293T cells were co-transfected with FLAG-tagged IRF5 and HA-tagged full-length or fragment eIF4G constructs. Cell lysates were subjected to immunoprecipitation and analyzed by immunoblotting as indicated. **C**, HEK293T cells were co-transfected with FLAG-DAP5 and HA-IRF5 and immunoprecipitated as indicated. Individual immunoprecipitation reactions were divided equally and incubated with or without 10 μg/mL RNase A at 37°C for 10 min prior to elution. **D**, IRF5(+) MEFs were incubated in complete media (CM) or EBSS for 2 h prior to cell lysis. Immunoprecipitation of endogenous DAP5, eIF4A, and PABP and analyzed by immunoblotting as indicated. DAP5 levels were below the limit of detection within input samples due to insufficient protein abundance relative to the percentage of input saved per reaction. **E**, IRF5(-) and IRF5(+) MEFs were incubated in complete media or EBSS for 2 h prior to cell lysis and lysates were subjected to m^7^GTP sepharose pull-down followed by immunoblotting as indicated. (**A-E**) All results are representative of n=3 independent experiments.

The eIF4G homologue, DAP5 (p97/eIF4G2) lacks the N-terminal eIF4E binding region that eIF4G possesses, but contains a MIF4G domain in similarity to one found in eIF4G (44,45). Therefore, we hypothesized that IRF5 may also interact with DAP5. Indeed, HA-tagged IRF5 co-immunoprecipitated with FLAG-tagged DAP5 **(Fig. 2c)**, further demonstrating that the interaction between IRF5 and the TIC is mediated through the MIF4G and independently of eIF4E.

Because the MIF4G domain also mediates interaction with RNA, we incubated the co-immunoprecipitation reactions with RNase A to exclude the possibility that our findings were the result of an indirect interaction between RNA and IRF5 (46). We observed that RNase A treatment had no effect on IRF5 interactions with DAP5 **(Fig. 2c)** or eIF4G **(Fig. S3a)**, indicating that the association between IRF5 and the MIF4G domain is the result of a protein-protein interaction. Similarly, RNase A treatment did not disrupt HA-tagged IRF5 co-immunoprecipitation of either FLAG-tagged eIF4A (**Fig. S3b**) or FLAG-tagged PABP (**Fig. S3c**) when co-expressed in HEK293T cells, suggesting that those associations are also the result of protein-protein interactions.

### IRF5-TIC interaction is associated with increased protein synthesis

To determine the functional effects of IRF5-TIC association, we used *Irf5*^*–/–*^ mouse embryonic fibroblasts (MEFs) to establish a genetically homogeneous cell line stably expressing HA-tagged human IRF5-V5 (denoted hereafter as IRF5(-) and IRF5(+), respectively) (47). IRF5-V5 shares 86% identity with the primary mouse IRF5 variant, and therefore, we expected that this would have no impact on the ability of IRF5 to interact with the TIC (48). To test whether we could re-capitulate IRF5-TIC interactions in MEF cells, we immunoprecipitated endogenous DAP5, eIF4A, and PABP from IRF5(+) cell lysates and measured interaction with HA-tagged IRF5. Western blot revealed that IRF5 interaction occurred between DAP5, eIF4A, and PABP in cells incubated in complete media (**Fig. 2d**). Because IRF5 interacted with the TIC independently of eIF4E, we also tested whether the IRF5 interaction with the TIC was sensitive to amino acid deprivation, a stress condition in which m^7^GTP-cap-independent mRNA translation is induced (49). We similarly demonstrated interaction between IRF5 and DAP5 as well as eIF4A, however we did not detect binding to PABP (**Fig. 2d**). We also tested the ability of HA-IRF5 to interact with eIF4E in MEFs by m^7^GTP pull-down. Consistent with our previous results obtained in human cell lines, we did not observe any interaction between HA-IRF5 and eIF4E in complete media or following incubation in Earle’s balanced salt solutions (EBSS) (**Fig. 2e**). These results suggest that IRF5 interacts with the TIC independently of eIF4E in MEFs in a manner similar to that observed in human cell lineages.

To determine the impact of IRF5 on global translational activity, we measured mRNA translation rates in MEFs cultured in complete media or EBSS (**Fig. 3a**) using a non-radioactive method of puromycin incorporation (50). Remarkably, IRF5(+) MEFs exhibited higher rates of mRNA translation compared to IRF5(-) MEFs in complete media (1.7-fold, *P* < 0.05) and following amino acid deprivation (4.8-fold, *P* < 0.0001) **(Fig. 3a, b)**. Amino acid deprivation with EBSS reduced protein synthesis in IRF5(-) MEFs an average of 2.9-fold (*P* < 0.05) versus no change in IRF5 (+) MEFs **(Fig. 3b)**. To examine whether structural variability in IRF5 exon 6 influences these changes in protein synthesis, we compared mRNA translation rates of IRF5(-) MEFs stably expressing either IRF5-V1, IRF5-V2/V6, or IRF5-V3/V4 to IRF5-V5 and IRF5(-) MEFs. Consistent with our earlier finding that IRF5 exon 6 structural variability has no impact on eIF4G interaction, we found that MEFs expressing IRF5 exon 6 structural variants yielded a comparable rate of mRNA translation to IRF5-V5 and a higher rate relative to IRF5 (-) MEFs **(Fig. S4a-c)**.

**FIGURE 3.**
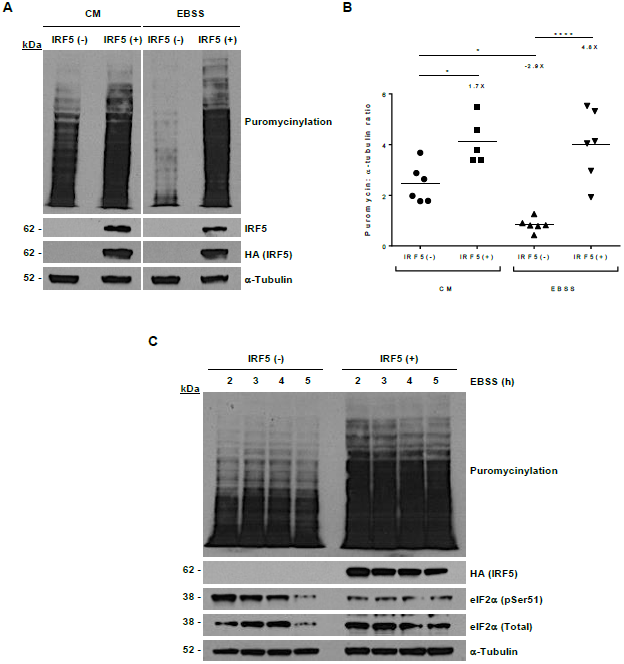
IRF5 increases the rate of mRNA translation. **A,** IRF5(-) and IRF5(+) MEFs were incubated in complete media (CM) or EBSS for a total of 2 h. At 1 hr prior to harvest, MEFs were incubated with 100 ug/mL puromycin, and then cell lysates were subjected to immunoblotting with a puromycin antibody. Data are representative of n=6 independent experiments. (**B**) Experiments were performed as in A and mRNA translation rates were quantified by the ratio of puromycin to α-tubulin band intensity. Data are representative of n=6 independent experiments. * *P* < 0.05, **** *P* < 0.0001 by Tukey’s multiple comparison test. (Note: 1 suspected outlier more than 3 standard deviations greater than the mean was identified by Grubbs’ test (α =0.05) in IRF5(+) CM-treated group and removed.) (**C**) IRF5(-) and IRF5(+) MEFs were cultured in EBSS as indicated, then mRNA translation rates were measured by puromycinylation as described in A. Results are representative of n=2 independent experiments.

In complete media, the slight increase in mRNA translation rate occurred without alteration in mTOR activity, as indicated by the similar levels of 4E-BP1 and RPS6 phosphorylation between IRF5(-) and IRF5(+) MEFs **(Fig. S4d)** (51). In the presence of nutrient deprivation, IRF5(+) MEFs sustained protein synthesis even in the absence of detectable phospho-4E-BP1 and phospho-RPS6, consistent with mTOR inhibition **(Fig. S4d)**. Since mTOR activity positively regulates eIF4E-cap-dependent translation via the reduction of 4E-BP1 affinity for eIF4E and thereby allowing eIF4E entry into the TIC, these data strongly suggest that mRNA translation rate in IRF5(+) MEFs is eIF4E-cap-independent (51).

Regulation of translation initiation can also occur independently of mTOR inhibition via phosphorylation of eIF2α Ser51 (p-eIF2α) by eIF2α kinases (52). To explore this mechanism, we measured the level of p-eIF2α in IRF5(-) and IRF5(+) MEFs from 2 hours to 5 hours following amino acid deprivation. Consistent with previous experiments, IRF5(+) MEFs demonstrated increased levels of protein synthesis compared to IRF5(-) MEFs at all time points evaluated (**Fig. 3c**). Furthermore, the increased protein synthesis observed in IRF5(+) MEFs was correlated with lower levels of p-eIF2α Therefore, the data suggest that IRF5-dependent changes in mRNA translation rate are associated with p-eIF2α status.

### IRF5 promotes mRNA translation of ATF4 and ATF4-dependent targets

Phosphorylation of eIF2α by the eIF2α kinase, GCN2, during the ISR results in translation of ATF4 mRNA transcripts followed by up-regulation of ATF4-dependent target genes including GADD34 (20). GADD34 is an eIF2α phosphatase that induces protein synthesis under stress (20,53). Therefore, we hypothesized that IRF5-dependent protein synthesis observed in **Fig. 3** and **Fig. S4** may be the result of ATF4 mRNA translation. We cultured IRF5(-) and IRF5(+) MEFs in EBSS for up to 12 hours and observed increased up-regulation of ATF4 as early as 15 minutes and as late as 4 hours post-EBSS in IRF5(+) MEFs (**Fig. 4a**). Peak levels of ATF4 were obtained at 1 hour post-EBSS, and subsequent experiments demonstrated a mean increase of 3.6-fold over IRF5(-) MEFs (*P* < 0.0001) (**Fig. 4b**). Using IRF5(+) MEF clones expressing different levels of HA-IRF5, we determined that EBSS-induced ATF4 protein synthesis was positively correlated with IRF5 expression **(Fig. S5)**. These data demonstrate that IRF5 up-regulates ATF4 during the ISR. These data indicate that the magnitude of ATF4 mRNA translation is responsive to IRF5 expression levels during the ISR.

**FIGURE 4.**
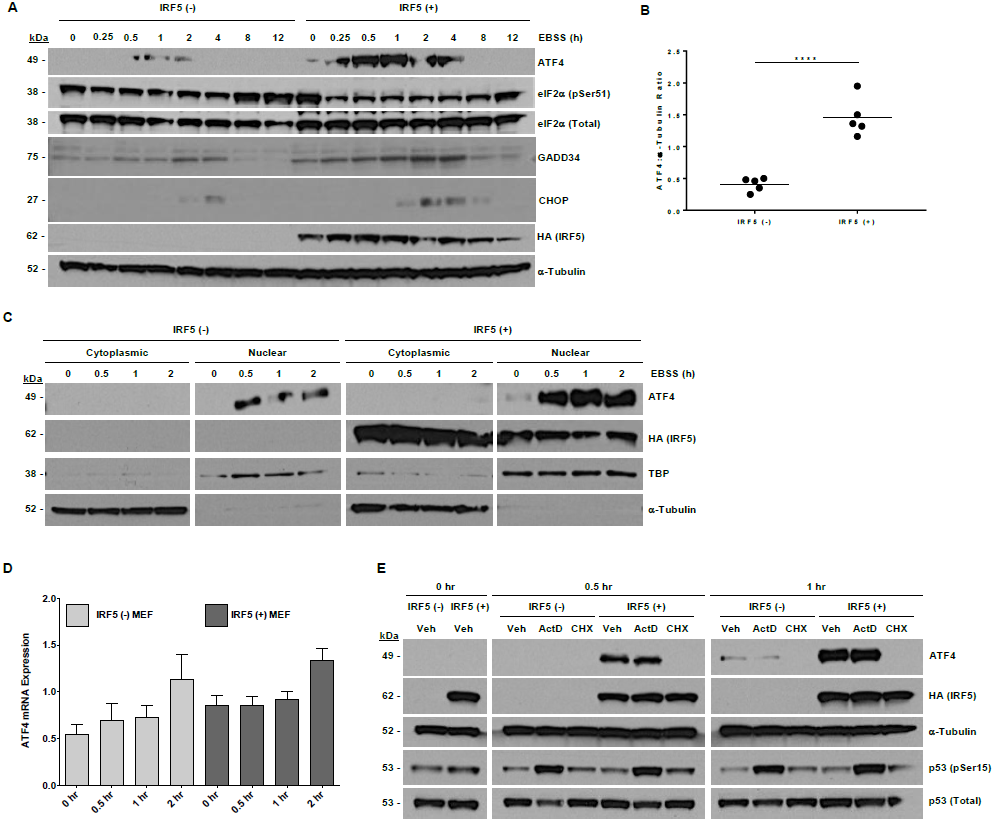
IRF5 promotes ATF4 mRNA translation and up-regulates ATF4-dependent target genes. **A,** IRF5(-) and IRF5(+) MEFs were cultured in EBSS as indicated and cell lysates were analyzed by immunoblotting. Results are representative of n=3 independent experiments. **B**, IRF5(-) and IRF5(+) MEFs were cultured in EBSS for 1 h and cell lysates were analyzed by immunoblotting. ATF4 protein expression was quantified the ratio of ATF4 to a-tubulin band intensity. Data are representative of n=3 independent experiments. **** *P* < 0.0001 by unpaired *t*-test. **C,** IRF5(-) and IRF5(+) MEFs were cultured in EBSS as indicated and cytoplasmic and nuclear lysates were analyzed by immunoblotting. Results are representative of n=3 independent experiments. **D**, Total mRNA was isolated from IRF5(-) and IRF5(+) MEFs cultured in EBSS for the indicated times. Samples were analyzed in triplicate by RT-PCR. *Atf4* mRNA expression was normalized to *Gapdh* and *Actb* and is displayed as mean ± SEM. Data is the result of n=3 independent experiments. **E,** IRF5(-) and IRF5(+) MEFs were either left untreated or pre-treated with DMSO, ActD (5 μg/mL), or CHX (50 μg/mL) for 1h, then cultured in EBSS for 0.5 h or 1 h supplemented with DMSO, ActD (5 μg/mL), or CHX (50 μg/mL). Immunoblotting of cell lysates was performed as indicated. Results are representative of n=3 independent experiments.

Additionally, expressed ATF4 was translocated to the nucleus suggestive of transcriptional activation (**Fig. 4c**). Consistent with up-regulation and nuclear localization of ATF4, we observed increased expression of the ATF4-dependent target genes, GADD34 and CHOP, in IRF5(+) MEFs (**Fig. 4a**). Furthermore, up-regulation of GADD34 correlated with decreased p-eIF2α similar to that shown in **Fig. 3c**. Unlike ATF4 expressed in MEFs, we did not observe increased IRF5 nuclear translocation following culture in EBSS (**Fig. 4c**). A similar result was obtained by immunofluorescence staining of IRF5(+) MEFs in response to EBSS where the majority of IRF5 remained concentrated in the cytoplasmic compartment **(Fig. S6)**. These data suggest that IRF5 is not activated by EBSS but is still able to up-regulate critical mediators of the ISR.

Since IRF5 is a transcription factor and a fraction is constitutively found within the nucleus (**Fig. 4c**), we evaluated whether IRF5 promoted transcription of the *Atf4* gene by culturing MEFs for up to 2 hours in EBSS. We did not observe significant changes in *Atf4* mRNA expression from baseline in either IRF5(-) or IRF5(+) MEFs (**Fig. 4d**). Nor did we observe any significant changes in *Atf4* mRNA expression between IRF5(-) and IRF5(+) MEFs at corresponding timepoints (**Fig. 4d**). These results suggest that IRF5 does not up-regulate ATF4 at the transcriptional level. To further explore this effect and determine whether IRF5 may regulate ATF4 expression at the level of mRNA translation, we pre-treated IRF5(-) and IRF5(+) MEFs in culture media for 1 hour with either an RNA synthesis inhibitor (actinomycin D; ActD), a protein synthesis inhibitor (cycloheximide; CHX), or DMSO (negative control) and then cultured cells for 30 minutes or 1 hour in EBSS containing fresh inhibitor. In agreement with data obtained in **Fig. 4a-c**, IRF5(+) MEFs treated with DMSO expressed higher levels of ATF4 compared to DMSO-treated IRF5(-) MEFs (**Fig. 4e**). Consistent with ATF4 regulation at the level of mRNA translation and the lack of IRF5 nuclear translocation, ATF4 protein synthesis in both IRF5(-) and IRF5(+) MEFs was unaffected by ActD treatment. However, ActD did inhibit transcription elongation as indicated by phosphorylation of p53 (**Fig. 4e**) (54). In contrast to ActD, treatment of IRF5(-) and IRF5(+) MEFs with CHX completely inhibited ATF4 expression in both cell lines (**Fig. 4e**). When considered together with our finding that IRF5 associates with the TIC, the data strongly suggest that IRF5 promotes ATF4 mRNA translation.

## DISCUSSION

Our study establishes a new role for IRF5 in regulating gene expression. The data demonstrate that IRF5 enhances expression of ISR-related genes following amino acid deprivation, and unlike previous reports of transcriptional control by IRF5, we determined that increased ATF4 up-regulation was at the level of mRNA translation (**Fig. 5**). Increased ATF4 expression in IRF5(+) MEFs resulted in higher levels of downstream ATF4-dependent targets, including GADD34 and CHOP, compared to those observed in IRF5(-) MEFs. Like ATF4, GADD34 and CHOP mRNA transcripts contain uORFs located within the 5’-leader region that negatively regulate translational efficiency under non-stress conditions, however activation of the ISR results in ribosomal bypass of uORFs and increased translation of the downstream coding region (55-57). While some aspects governing this process are known, the precise regulatory mechanisms are not defined. In our study, we detected ATF4, GADD34, and CHOP protein expression at earlier time points in IRF5(+) MEFs following amino acid deprivation. Additionally, baseline ATF4 and GADD34 levels were also higher compared to those observed in IRF5(-) MEFs. Therefore, our data may suggest that IRF5 facilitates ribosomal recruitment to alternative translation start sites such as those found in ATF4, GADD34, and CHOP thereby increasing their protein synthesis and conveying increased cell sensitivity to changes in nutrient levels.

**FIGURE 5.**
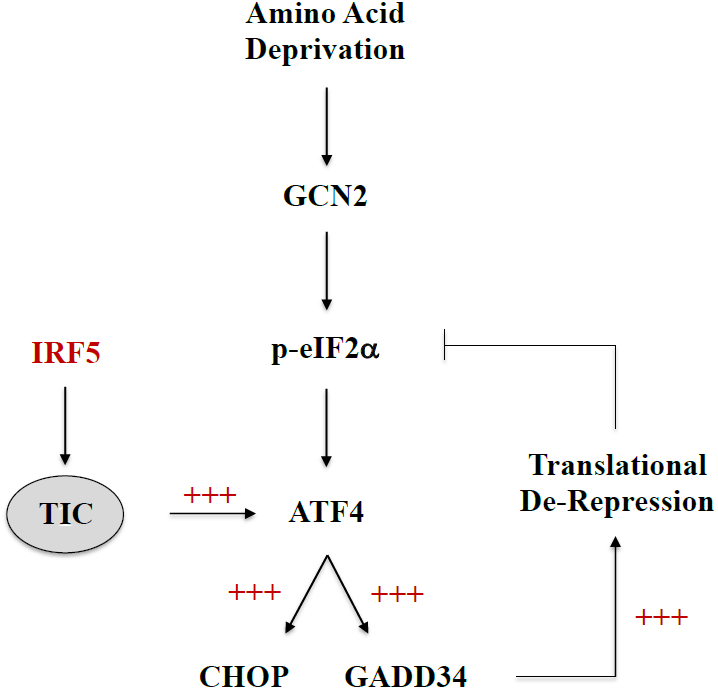
Model of IRF5-dependent regulation of the ISR. Uncharged tRNAs bind to GCN2 inducing its activation during amino acid deprivation. GCN2 then phosphorylates eIF2α leading to a slower rate of ternary complex recruitment to the TIC and subsequent ribosomal bypass of the negative regulatory element in the ATF4 5’ leader region. IRF5 association with the TIC may facilitate this process resulting in enhanced ATF4 mRNA translation. Increased ATF4 up-regulates expression of the ATF4-dependent target genes, CHOP and GADD34. Higher levels of GADD34 augments the negative feedback loop that promotes translational de-repression during the ISR.

Along these lines, we showed that IRF5 associates with the TIC including the scaffolding proteins, eIF4G and DAP5. Furthermore, we demonstrated the importance of the MIF4G domain for facilitating this interaction. The MIF4G domain contains binding sites for the eIF3 complex as well as eIF4A, which we also established as an interaction partner of IRF5 (44). The eIF3 complex is responsible for 40S ribosomal recruitment and is a structure homologous to the COP9 signalosome (CSN) (58). Interestingly, an interaction between IRF5 and CSN was described in a previous report (59). Our findings may indicate that IRF5 possesses affinity for structures bearing similarity to CSN (e.g. eIF3) and may interact with the TIC via this complex.

During our effort to define IRF5-TIC interactions, we were unable to establish any association between IRF5 and the m^7^GTP-binding protein, eIF4E, which promotes cap-dependent mRNA translation (1,4). However, we cannot rule out the possibility that IRF5 may interact with other cap-binding proteins instead of eIF4E. Additionally, 4E-BP1-mediated repression of eIF4E-eIF4G interaction does not appear to explain incorporation of IRF5 into the TIC, since we observed IRF5-eIF4G interaction following culture in complete media when 4E-BP-1 affinity for eIF4E is lowest (60). Although we demonstrated that IRF5 interacts with DAP5 which does not contain an eIF4E binding region, we cannot eliminate the possibility that IRF5 binding to the MIF4G domain of eIF4G interferes with eIF4E binding.

Cap-dependent mRNA translation is the predominant mode of protein synthesis used for a variety of cell functions including growth and proliferation (61). However, different forms of cell stress trigger shut-down of this process and cap-independent mRNA translation then enables up-regulation of specific genes important for mediating the cell stress response (62). In addition to alternative ORFs, certain mRNA transcripts reportedly contain cis-regulatory elements known as internal ribosomal entry sites (IRESs) that are regulated by transacting factors which promote translation initiation of stress response genes (62). In our study, we detected multiple putative IRES transacting factors by IP-mass spectrometry in association with IRF5 including Splicing Factor Proline-Glutamine Rich (SFPQ), Interleukin Enhancer Binding Factor 2 (ILF2), and members of the heterogenous nuclear ribonucleoprotein (hnRNP) family (63-68). Coupled with the lack of interaction observed between IRF5 and eIF4E, this may suggest that IRF5 regulates mRNA translation via cap-independent mechanisms.

IRF5 over-expression is associated with multiple autoimmune diseases including systemic lupus erythematosus (SLE) and systemic sclerosis (SSc) (69,70). The role for IRF5 in disease pathogenesis is often associated with the increased production of type I interferon and pro-inflammatory cytokines (71-73). While dysregulation of type I interferon and cytokines frequently accompany these conditions, the true etiology is unknown. These complex diseases are multifactorial and the result of both genetic and environmental factors. Our finding that IRF5 regulates the ISR in response to amino acid deprivation requires further investigation within the context of human disease and will take into consideration other forms of cell stress that trigger the ISR such as the accumulation of unfolded proteins and dsRNA which activate PERK and PKR, respectively (14). These studies may provide new insight into the significance of changes in GADD34 expression and CHOP activation that have been reported in autoimmune disease patient cells.

## EXPERIMENTAL PROCEDURES

### Reagents and Antibodies

Cycloheximide (50 μL/mL; 239765) and nuclear extraction kits (2900) were purchased from EMD Millipore; Actinomycin D (5 μg/mL; A1410), RNase A (10 μg/mL; R4642), and DMSO (D2438) were purchased from Sigma; Cell culture grade DMSO was purchased from Sigma-Aldrich. Antibodies against FLAG (8146S), HA (3724S), 4E-BP1 (9644S), 4E-BP1 pSer65 (9451S), RPS6 (2217S), RPS6 pSer235-236 (4857S), eIF2α (2103S), ATF4 (11815S), CHOP (28955), p53 pSer15 (12571P), and p53 (2524S) were all purchased from Cell Signaling Technology; Antibodies against eIF2α pSer51 (ab32157), eIF4A1 (ab31217), eIF4E (ab1126), IRF5 (ab21689), IRF5 (ab2932), PABP (ab21060), α-Tubulin (ab7291), and TATA-binding protein (TBP; ab818) were from Abcam; Antibodies against eIF4G1 (RN002P) and eIF4G2/DAP5 (RN003P) were from MBL International; Antibodies against GADD34 (10087-304) were from Protein Tech; Purified rabbit IgG (12-370 was from Millipore; Normal Rabbit Serum (ab166640) was from Abcam. Horseradish Peroxidase (HRP)-conjugated mouse IgG (HAF018), rabbit IgG (HAF008) or goat IgG (HAF109) were purchased from R&D.

### Cell Culture

HEK293T, THP1, and BJAB cells were originally obtained from ATCC (CRL-3216, TIB-202). IRF5^-/-^ MEF cells were originally obtained from Dr. Tak Mak, University of Toronto, Ontario, Canada, and developed as described (47,74,75). MEF and HEK293T cells were maintained in DMEM; BJAB and THP1 cells were maintained in RPMI. All cell lines were supplemented with 10% FBS (Atlanta Biologicals; 501520HI), 2mM L-glutamine (VWR; 12001-698), and 50 U/mL Penicillin-Streptomycin (Fischer Scientific; 15070-063); these media are herein referred to as complete medium. Earle’s Balanced Salt Solution (EBSS) (Life Technologies; 14155-063) was used as nutrient deprivation media. For experiments using EBSS starvation, MEFs were first plated at 5×10^5^ cells/mL into a 10-cm dish in complete medium and cultured for 16-19 hours or until 80-85% confluence. Cells were washed twice with pre-warmed PBS before culturing in EBSS.

### Expression Vectors and Retroviral Gene Transfer

cDNAs of human IRF5 variants, V5, V1, V2/V6, and V3/V4, were developed and used for retroviral gene transfer to create HA-tagged or FLAG-tagged IRF5 V5, V1, V2/V6 or V3/V4 MEFs as previously described (76). N-terminal HA-tagged eIF4G1 plasmids were generated using pcDNA3 vector as previously described (77). FLAG-eIF4A and FLAG-PABP expression vectors were obtained from GenScript. Detailed information is provided in the Supplemental Experimental Procedures.

### Whole Cell Lysates and Immunoblotting

Cytoplasmic & nuclear extracts were generated via a nuclear extraction kit (EMD Millipore, 2900) according to protocol. Cells or cell extracts were washed in cold PBS and lysed with RIPA or NP40 lysis buffer (20 mM Tris-HCl, 137 mM NaCl, 2 mM EDTA, pH 8, 10% glycerol, 1% NP40) containing protease and Halt phosphatase inhibitors (EMD Millipore and Pierce, respectively). Protein concentrations were determined by Qubit assay (Life Technologies). Proteins were denatured in 2X SDS loading buffer and heated to 100°C for 5-10 min. Proteins were eluted by SDS-PAGE and analyzed by immunoblotting as indicated.

### Co-Immunoprecipitation

To co-immunoprecipitate endogenous proteins, cells were first cross-linked with 1% formaldehyde (final volume) for 5 min at room temperature. Formaldehyde was then neutralized with 0.125 M glycine (final volume) and cell pellets washed twice with cold PBS prior to lysis with NP40 lysis buffer described above. Experiments using HA- and FLAG-tagged proteins were performed as described above without cross-linking. Protein concentrations were determined by Qubit assay and used to balance loading of immunoprecipitation reactions; a total of 1000 μg of cell lysate was brought to a volume of 500 μL in NP40 lysis buffer. Lysates were pre-cleared with 30 μL of protein A/G magnetic beads (EMD Millipore) prior to immunoprecipitation with antibody-coated protein A/G magnetic beads. Reactions were incubated overnight with rotation at 4°C and then washed five times with NP40 lysis buffer followed by elution with 2X SDS loading buffer. Samples were heated to 100°C for 5 minutes followed by SDS-PAGE. Antibodies to eIF4G1 (RN002P) and eIF4G2 (RN003P) were from MBL International and were used at 15 μg/IP; anti-eIF4A1 (ab31217), anti-PABP (ab21060), and anti-IRF5 (ab2932) were from Abcam and used at 15 μg/IP; anti-eIF4E (ab1126) was also from Abcam but used at 3 μL/IP; anti-FLAG (8146S) and anti-HA (3724S) were from Cell Signaling Technology and were used at 3 μg/IP; purified rabbit IgG (millipore) or normal rabbit serum (Abcam) were used as isotype controls in an amount equivalent to the target-specific antibody.

### Affinity Purification and Liquid Chromatography-Mass Spectrometry Analysis

Co-immunoprecipitation and protein elution were performed as described above. Proteins were resolved, then processed for and analyzed by LC-Mass Spectroscopy as previously described (78). Detailed information is provided in the Supplemental Experimental Procedures.

### 7-methyl GTP-Sepharose Binding Assay

*7***-**methyl GTP (m^7^GTP) sepharose (Jena Bioscience) and sepharose (Sigma) resins were blocked by pre-incubating in 0.2M glycine with rotation at 4°C for 1 h. Cells were lysed in NP40 lysis buffer and protein concentrations quantified by Qubit assay. A total of 500 μg of protein and 60-μL volume of m^7^GTP sepharose or sepharose resin were brought to a final volume of 500 μL with binding buffer (50 mM Tris-HCl, 30 mM NaCl, 2.5 mM MgCl2, pH 7.5, 5% glycerol, 0.1% 2-mercaptoethanol). Pull-down reactions were incubated overnight with rotation at 4°C. Reactions were washed five times with binding buffer and eluted with 2X SDS loading buffer by boiling for 5-10 min. Eluted proteins were resolved by SDS-PAGE and immunoblotting.

### SUnSET Assay

SUnSET Assay was used to puromycinylate translated proteins as previously described (50).

### Immunofluorescence Microscopy

MEFs were cultured on cover slips at 37°C. Cells were fixed in 4% paraformaldehyde for 10 min followed by three rinses with PBS. Cells were permeabilized with 0.1% Tween-20 and rinsed with PBS prior to blocking with 5% normal donkey serum for 20 min. Primary antibodies to monoclonal rabbit anti-HA (Cell Signaling Technology; 3724S) and monoclonal mouse anti-5.8S rRNA (Life Technologies; MA1-13017) were incubated overnight at 4°C in a humidified chamber at a concentration of 1:3200 and 1:100, respectively. The next day slides were rinsed and incubated for 45 min at room temperature with Alexa Fluor 488-conjugated donkey anti-rabbit and Cy3-conjugated donkey anti-mouse secondary antibodies (Jackson ImmunoResearch Laboratories; 711-546-152, 705-167-003) at a concentration of 1:2400 and 1:1200, respectively. Cells were rinsed with PBS and stained with DAPI (1:600) for 2 min. Cells were rinsed a final time with PBS and cover-slipped prior to imaging with a Zeiss LSM710 confocal laser-scanning microscope.

### Gene expression analysis by RT-PCR

Cells were treated as described and total RNA extracted using an RNA Clean and Concentrator kit-25 (Zymo Research; R1017) according to manufacturer’s instruction to purify total RNA. RNA concentrations were measured by Qubit BR assay (Life Technologies; Q10211) and evenly loaded into reverse transcription (RT) reactions for conversion to cDNA using an iScript cDNA Synthesis Kit (Bio-Rad; 170-8891). Equal volumes of each sample were loaded into RT-PCR reactions that were performed in triplicate using RT^2^ SYBR Green qPCR master mix (Qiagen; 330502) and validated primer sets to target genes (Integrated DNA Technologies); primer sequences are available upon request. Relative ATF4 expression was determined via the ΔΔCT method by normalization to β-actin and GAPDH using CFX Manager Software (Bio-Rad).

### Statistical analysis

All quantified data are presented as the mean ± SEM of at least five independent experiments. GraphPad Prism (V7.02) software was used for statistical analysis. Statistical test implemented and resulting *P*-values are as indicated. Grubbs’ test was used for outlier detection.

## Acknowledgements

We thank J. Maiers and the OMRF Imaging Core Facility for their assistance with immunofluorescence microscopy. We also thank N. Sonenberg for his critical review of this manuscript. This work was supported by grants from the National Institutes of Health (R01 AR063124, R01 AR056360, U19 AI082714, P30 GM110766).

## Conflicts of interest

The authors declare no competing financial interests.

## Author contributions

Conceptualization, J.D.W. and P.M.G; Methodology, Investigations and Validation, J.D.W., M.W., Y.W., M.K., F.W. and, A.Y.; Formal Analysis, J.D.W.; Resources, N.S.; Writing – Original Draft, J.D.W. and P.M.G; Writing – Review & Editing, J.D.W., K.L.T, and P.M.G.; Supervision, Project Administration, and Funding Acquisition, P.M.G.

## REFERENCES

1. Sonenberg, N., Rupprecht, K. M., Hecht, S. M., and Shatkin, A. J. (1979) Eukaryotic mRNA cap binding protein: purification by affinity chromatography on sepharose-coupled m7GDP. Proceedings of the National Academy of Sciences of the United States of America 76, 4345–4349

2. Ray, B. K., Lawson, T. G., Kramer, J. C., Cladaras, M. H., Grifo, J. A., Abramson, R. D., Merrick, W. C., and Thach, R. E. (1985) ATP-dependent unwinding of messenger RNA structure by eukaryotic initiation factors. Journal of Biological Chemistry 260, 7651–7658

3. Lamphear, B. J., Kirchweger, R., Skern, T., and Rhoads, R. E. (1995) Mapping of Functional Domains in Eukaryotic Protein Synthesis Initiation Factor 4G (eIF4G) with Picornaviral Proteases: IMPLICATIONS FOR CAP-DEPENDENT AND CAP-INDEPENDENT TRANSLATIONAL INITIATION. Journal of Biological Chemistry 270, 21975–21983

4. Sonenberg, N., Morgan, M. A., Merrick, W. C., and Shatkin, A. J. (1978) A polypeptide in eukaryotic initiation factors that crosslinks specifically to the 5’-terminal cap in mRNA. Proceedings of the National Academy of Sciences of the United States of America 75, 4843–4847

5. Trachsel, H., and Staehelin, T. (1979) Initiation of mammalian protein synthesis. The multiple functions of the initiation factor eIF-3. Biochimica et biophysica acta 565, 305–314

6. Benne, R., and Hershey, J. W. (1978) The mechanism of action of protein synthesis initiation factors from rabbit reticulocytes. Journal of Biological Chemistry 253, 3078–3087

7. Raychaudhuri, P., Chaudhuri, A., and Maitra, U. (1985) Formation and release of eukaryotic initiation factor 2 X GDP complex during eukaryotic ribosomal polypeptide chain initiation complex formation. Journal of Biological Chemistry 260, 2140–2145

8. Konieczny, A., and Safer, B. (1983) Purification of the eukaryotic initiation factor 2-eukaryotic initiation factor 2B complex and characterization of its guanine nucleotide exchange activity during protein synthesis initiation. The Journal of Biological Chemistry 258, 3402–3408

9. Donnelly, N., Gorman, A. M., Gupta, S., and Samali, A. (2013) The eIF2alpha kinases: their structures and functions. Cellular and molecular life sciences: CMLS 70, 3493–3511

10. Taniuchi, S., Miyake, M., Tsugawa, K., Oyadomari, M., and Oyadomari, S. (2016) Integrated stress response of vertebrates is regulated by four eIF2α kinases. 6, 32886

11. Berlanga, J. J., Ventoso, I., Harding, H. P., Deng, J., Ron, D., Sonenberg, N., Carrasco, L., and de Haro, C. (2006) Antiviral effect of the mammalian translation initiation factor 2alpha kinase GCN2 against RNA viruses. The EMBO journal 25, 1730–1740

12. del Pino, J., Jiménez, J. L., Ventoso, I., Castelló, A., Muñoz-Fernández, M. á., de Haro, C., and Berlanga, J. J. (2012) GCN2 Has Inhibitory Effect on Human Immunodeficiency Virus-1 Protein Synthesis and Is Cleaved upon Viral Infection. PLoS ONE 7, e47272

13. Wek, S. A., Zhu, S., and Wek, R. C. (1995) The histidyl-tRNA synthetase-related sequence in the eIF-2 alpha protein kinase GCN2 interacts with tRNA and is required for activation in response to starvation for different amino acids. Mol Cell Biol 15, 4497–4506

14. Pakos-Zebrucka, K., Koryga, I., Mnich, K., Ljujic, M., Samali, A., and Gorman, A. M. (2016) The integrated stress response. EMBO reports 17, 1374–1395

15. Harding, H. P., Novoa, I., Zhang, Y., Zeng, H., Wek, R., Schapira, M., and Ron, D. (2000) Regulated translation initiation controls stress-induced gene expression in mammalian cells. Molecular cell 6, 1099–1108

16. Harding, H. P., Zhang, Y., Zeng, H., Novoa, I., Lu, P. D., Calfon, M., Sadri, N., Yun, C., Popko, B., Paules, R., Stojdl, D. F., Bell, J. C., Hettmann, T., Leiden, J. M., and Ron, D. (2003) An integrated stress response regulates amino acid metabolism and resistance to oxidative stress. Molecular cell 11, 619–633

17. Siu, F., Bain, P. J., LeBlanc-Chaffin, R., Chen, H., and Kilberg, M. S. (2002) ATF4 is a mediator of the nutrient-sensing response pathway that activates the human asparagine synthetase gene. The Journal of Biological Chemistry 277, 24120–24127

18. Ohoka, N., Yoshii, S., Hattori, T., Onozaki, K., and Hayashi, H. (2005) TRB3, a novel ER stress-inducible gene, is induced via ATF4-CHOP pathway and is involved in cell death. The EMBO journal 24, 1243–1255

19. B’ Chir, W., Maurin, A. C., Carraro, V., Averous, J., Jousse, C., Muranishi, Y., Parry, L., Stepien, G., Fafournoux, P., and Bruhat, A. (2013) The eIF2alpha/ATF4 pathway is essential for stress-induced autophagy gene expression. Nucleic acids research 41, 7683–7699

20. Ma, Y., and Hendershot, L. M. (2003) Delineation of a Negative Feedback Regulatory Loop That Controls Protein Translation during Endoplasmic Reticulum Stress. Journal of Biological Chemistry 278, 34864–34873

21. Vattem, K. M., and Wek, R. C. (2004) Reinitiation involving upstream ORFs regulates ATF4 mRNA translation in mammalian cells. Proceedings of the National Academy of Sciences of the United States of America 101, 11269–11274

22. Proenca-Modena, J. L., Hyde, J. L., Sesti-Costa, R., Lucas, T., Pinto, A. K., Richner, J. M., Gorman, M. J., Lazear, H. M., and Diamond, M. S. (2016) Interferon-Regulatory Factor 5-Dependent Signaling Restricts Orthobunyavirus Dissemination to the Central Nervous System. Journal of Virology 90, 189–205

23. Thackray, L. B., Shrestha, B., Richner, J. M., Miner, J. J., Pinto, A. K., Lazear, H. M., Gale, M., and Diamond, M. S. (2014) Interferon Regulatory Factor 5-Dependent Immune Responses in the Draining Lymph Node Protect against West Nile Virus Infection. Journal of Virology 88, 11007–11021

24. Hu, G., and Barnes, B. J. (2009) IRF-5 is a mediator of the death receptor-induced apoptotic signaling pathway. The Journal of Biological Chemistry 284, 2767–2777

25. Couzinet, A., Tamura, K., Chen, H. M., Nishimura, K., Wang, Z., Morishita, Y., Takeda, K., Yagita, H., Yanai, H., Taniguchi, T., and Tamura, T. (2008) A cell-type-specific requirement for IFN regulatory factor 5 (IRF5) in Fas-induced apoptosis. Proceedings of the National Academy of Sciences of the United States of America 105, 2556–2561

26. Hu, G., Mancl, M. E., and Barnes, B. J. (2005) Signaling through IFN regulatory factor-5 sensitizes p53-deficient tumors to DNA damage-induced apoptosis and cell death. Cancer research 65, 7403–7412

27. Schoenemeyer, A., Barnes, B. J., Mancl, M. E., Latz, E., Goutagny, N., Pitha, P. M., Fitzgerald, K. A., and Golenbock, D. T. (2005) The interferon regulatory factor, IRF5, is a central mediator of toll-like receptor 7 signaling. The Journal of Biological Chemistry 280, 17005–17012

28. Steinhagen, F., McFarland, A. P., Rodriguez, L. G., Tewary, P., Jarret, A., Savan, R., and Klinman, D. M. (2013) IRF-5 and NF-kappaB p50 co-regulate IFN-beta and IL-6 expression in TLR9-stimulated human plasmacytoid dendritic cells. European journal of immunology 43, 1896–1906

29. Feng, D., Sangster-Guity, N., Stone, R., Korczeniewska, J., Mancl, M. E., Fitzgerald-Bocarsly, P., and Barnes, B. J. (2010) Differential requirement of histone acetylase and deacetylase activities for IRF5-mediated proinflammatory cytokine expression. Journal of immunology (Baltimore, Md.: 1950) 185, 6003–6012

30. Barnes, B. J., Kellum, M. J., Pinder, K. E., Frisancho, J. A., and Pitha, P. M. (2003) Interferon regulatory factor 5, a novel mediator of cell cycle arrest and cell death. Cancer research 63, 6424–6431

31. Pimenta, E. M., and Barnes, B. J. (2015) A conserved region within interferon regulatory factor 5 controls breast cancer cell migration through a cytoplasmic and transcription-independent mechanism. Mol Cancer 14, 32

32. Aitken, C. E., and Lorsch, J. R. (2012) A mechanistic overview of translation initiation in eukaryotes. Nat Struct Mol Biol 19, 568–576

33. Komar, A. A., and Hatzoglou, M. (2011) Cellular IRES-mediated translation: the war of ITAFs in pathophysiological states. Cell Cycle 10, 229–240

34. Parsyan, A., Svitkin, Y., Shahbazian, D., Gkogkas, C., Lasko, P., Merrick, W. C., and Sonenberg, N. (2011) mRNA helicases: the tacticians of translational control. Nat Rev Mol Cell Biol 12, 235–245

35. Mancl, M. E., Hu, G., Sangster-Guity, N., Olshalsky, S. L., Hoops, K., Fitzgerald-Bocarsly, P., Pitha, P. M., Pinder, K., and Barnes, B. J. (2005) Two discrete promoters regulate the alternatively spliced human interferon regulatory factor-5 isoforms. Multiple isoforms with distinct cell type-specific expression, localization, regulation, and function. The Journal of biological chemistry 280, 21078–21090

36. Stone, R. C., Du, P., Feng, D., Dhawan, K., Ronnblom, L., Eloranta, M. L., Donnelly, R., and Barnes, B. J. (2013) RNA-Seq for enrichment and analysis of IRF5 transcript expression in SLE. PLoS ONE 8, e54487

37. Li, W., Belsham, G. J., and Proud, C. G. (2001) Eukaryotic initiation factors 4A (eIF4A) and 4G (eIF4G) mutually interact in a 1:1 ratio in vivo. The Journal of biological chemistry 276, 29111–29115

38. Safaee, N., Kozlov, G., Noronha, A. M., Xie, J., Wilds, C. J., and Gehring, K. (2012) Interdomain allostery promotes assembly of the poly(A) mRNA complex with PABP and eIF4G. Molecular cell 48, 375–386

39. Imataka, H., and Sonenberg, N. (1997) Human eukaryotic translation initiation factor 4G (eIF4G) possesses two separate and independent binding sites for eIF4A. Molecular and cellular biology 17, 6940–6947

40. Lamphear, B. J., Kirchweger, R., Skern, T., and Rhoads, R. E. (1995) Mapping of functional domains in eukaryotic protein synthesis initiation factor 4G (eIF4G) with picornaviral proteases. Implications for cap-dependent and cap-independent translational initiation. J Biol Chem 270, 21975–21983

41. Le, H., Tanguay, R. L., Balasta, M. L., Wei, C. C., Browning, K. S., Metz, A. M., Goss, D. J., and Gallie, D. R. (1997) Translation initiation factors eIF-iso4G and eIF-4B interact with the poly(A)-binding protein and increase its RNA binding activity. The Journal of biological chemistry 272, 16247–16255

42. Mader, S., Lee, H., Pause, A., and Sonenberg, N. (1995) The translation initiation factor eIF-4E binds to a common motif shared by the translation factor eIF-4 gamma and the translational repressors 4E-binding proteins. Molecular and cellular biology 15, 4990–4997

43. Pyronnet, S., Imataka, H., Gingras, A. C., Fukunaga, R., Hunter, T., and Sonenberg, N. (1999) Human eukaryotic translation initiation factor 4G (eIF4G) recruits mnk1 to phosphorylate eIF4E. The EMBO journal 18, 270–279

44. Virgili, G., Frank, F., Feoktistova, K., Sawicki, M., Sonenberg, N., Fraser, C. S., and Nagar, B. (2013) Structural analysis of the DAP5 MIF4G domain and its interaction with eIF4A. Structure 21, 517–527

45. Imataka, H., Olsen, H. S., and Sonenberg, N. (1997) A new translational regulator with homology to eukaryotic translation initiation factor 4G. The EMBO journal 16, 817–825

46. Pestova, T. V., Shatsky, I. N., and Hellen, C. U. (1996) Functional dissection of eukaryotic initiation factor 4F: the 4A subunit and the central domain of the 4G subunit are sufficient to mediate internal entry of 43S preinitiation complexes. Molecular and cellular biology 16, 6870–6878

47. Takaoka, A., Yanai, H., Kondo, S., Duncan, G., Negishi, H., Mizutani, T., Kano, S., Honda, K., Ohba, Y., Mak, T. W., and Taniguchi, T. (2005) Integral role of IRF-5 in the gene induction programme activated by Toll-like receptors. Nature 434, 243–249

48. Paun, A., Reinert, J. T., Jiang, Z., Medin, C., Balkhi, M. Y., Fitzgerald, K. A., and Pitha, P. M. (2008) Functional characterization of murine interferon regulatory factor 5 (IRF-5) and its role in the innate antiviral response. The Journal of biological chemistry 283, 14295–14308

49. Merrick, W. C. (2004) Cap-dependent and cap-independent translation in eukaryotic systems. Gene 332, 1–11

50. Schmidt, E. K., Clavarino, G., Ceppi, M., and Pierre, P. (2009) SUnSET, a nonradioactive method to monitor protein synthesis. Nat Methods 6, 275–277

51. Dowling, R. J., Topisirovic, I., Fonseca, B. D., and Sonenberg, N. (2010) Dissecting the role of mTOR: lessons from mTOR inhibitors. Biochimica et biophysica acta 1804, 433–439

52. Jackson, R. J., Hellen, C. U., and Pestova, T. V. (2010) The mechanism of eukaryotic translation initiation and principles of its regulation. Nat Rev Mol Cell Biol 11, 113–127

53. Novoa, I., Zeng, H., Harding, H. P., and Ron, D. (2001) Feedback inhibition of the unfolded protein response by GADD34-mediated dephosphorylation of eIF2alpha. The Journal of cell biology 153, 1011–1022

54. Choong, M. L., Yang, H., Lee, M. A., and Lane, D. P. (2009) Specific activation of the p53 pathway by low dose actinomycin D: a new route to p53 based cyclotherapy. Cell cycle (Georgetown, Tex.) 8, 2810–2818

55. Palam, L. R., Baird, T. D., and Wek, R. C. (2011) Phosphorylation of eIF2 Facilitates Ribosomal Bypass of an Inhibitory Upstream ORF to Enhance CHOP Translation. The Journal of Biological Chemistry 286, 10939–10949

56. Lee, Y.-Y., Cevallos, R. C., and Jan, E. (2009) An Upstream Open Reading Frame Regulates Translation of GADD34 during Cellular Stresses That Induce eIF2α Phosphorylation. The Journal of Biological Chemistry 284, 6661–6673

57. Jousse, C., Bruhat, A., Carraro, V., Urano, F., Ferrara, M., Ron, D., and Fafournoux, P. (2001) Inhibition of CHOP translation by a peptide encoded by an open reading frame localized in the chop 5’ UTR. Nucleic acids research 29, 4341–4351

58. Enchev, R. I., Schreiber, A., Beuron, F., and Morris, E. P. (2010) Structural insights into the COP9 signalosome and its common architecture with the 26S proteasome lid and eIF3. Structure 18, 518–527

59. Korczeniewska, J., and Barnes, B. J. (2013) The COP9 Signalosome Interacts with and Regulates Interferon Regulatory Factor 5 Protein Stability. Molecular and Cellular Biology 33, 1124–1138

60. Gingras, A. C., Gygi, S. P., Raught, B., Polakiewicz, R. D., Abraham, R. T., Hoekstra, M. F., Aebersold, R., and Sonenberg, N. (1999) Regulation of 4E-BP1 phosphorylation: a novel two-step mechanism. Genes & development 13, 1422–1437

61. Richter, J. D., and Sonenberg, N. (2005) Regulation of cap-dependent translation by eIF4E inhibitory proteins. Nature 433, 477–480

62. Hinnebusch, A. G., Ivanov, I. P., and Sonenberg, N. (2016) Translational control by 5′-untranslated regions of eukaryotic mRNAs. Science (New York, N.Y.) 352, 1413–1416

63. Graber, T. E., Baird, S. D., Kao, P. N., Mathews, M. B., and Holcik, M. (2010) NF45 functions as an IRES trans-acting factor that is required for translation of cIAP1 during the unfolded protein response. Cell death and differentiation 17, 719–729

64. Bonnal, S., Pileur, F., Orsini, C., Parker, F., Pujol, F., Prats, A. C., and Vagner, S. (2005) Heterogeneous nuclear ribonucleoprotein A1 is a novel internal ribosome entry site trans-acting factor that modulates alternative initiation of translation of the fibroblast growth factor 2 mRNA. The Journal of Biological Chemistry 280, 4144–4153

65. Kim, J. H., Paek, K. Y., Choi, K., Kim, T. D., Hahm, B., Kim, K. T., and Jang, S. K. (2003) Heterogeneous nuclear ribonucleoprotein C modulates translation of c-myc mRNA in a cell cycle phase-dependent manner. Mol Cell Biol 23, 708–720

66. Holcik, M., Gordon, B. W., and Korneluk, R. G. (2003) The internal ribosome entry site-mediated translation of antiapoptotic protein XIAP is modulated by the heterogeneous nuclear ribonucleoproteins C1 and C2. Mol Cell Biol 23, 280–288

67. Cammas, A., Pileur, F., Bonnal, S., Lewis, S. M., Leveque, N., Holcik, M., and Vagner, S. (2007) Cytoplasmic relocalization of heterogeneous nuclear ribonucleoprotein A1 controls translation initiation of specific mRNAs. Molecular biology of the cell 18, 5048–5059

68. Sharathchandra, A., Lal, R., Khan, D., and Das, S. (2012) Annexin A2 and PSF proteins interact with p53 IRES and regulate translation of p53 mRNA. RNA Biology 9, 1429–1439

69. Sigurdsson, S., Goring, H. H., Kristjansdottir, G., Milani, L., Nordmark, G., Sandling, J. K., Eloranta, M. L., Feng, D., Sangster-Guity, N., Gunnarsson, I., Svenungsson, E., Sturfelt, G., Jonsen, A., Truedsson, L., Barnes, B. J., Alm, G., Ronnblom, L., and Syvanen, A. C. (2008) Comprehensive evaluation of the genetic variants of interferon regulatory factor 5 (IRF5) reveals a novel 5 bp length polymorphism as strong risk factor for systemic lupus erythematosus. Human molecular genetics 17, 872–881

70. Carmona, F. D., Martin, J.-E., Beretta, L., Simeón, C. P., Carreira, P. E., Callejas, J. L., Fernández-Castro, M., Sáez-Comet, L., Beltrán, E., Camps, M. T., Egurbide, M. V., the Spanish Scleroderma, G., Airó, P., Scorza, R., Lunardi, C., Hunzelmann, N., Riemekasten, G., Witte, T., Kreuter, A., Distler, J. H. W., Madhok, R., Shiels, P., van Laar, J. M., Fonseca, C., Denton, C., Herrick, A., Worthington, J., Schuerwegh, A. J., Vonk, M. C., Voskuyl, A. E., Radstake, T. R. D. J., and Martín, J. (2013) The Systemic Lupus Erythematosus IRF5 Risk Haplotype Is Associated with Systemic Sclerosis. PLoS ONE 8, e54419

71. Niewold, T. B., Kelly, J. A., Flesch, M. H., Espinoza, L. R., Harley, J. B., and Crow, M. K. (2008) Association of the IRF5 risk haplotype with high serum interferon-alpha activity in systemic lupus erythematosus patients. Arthritis and rheumatism 58, 2481–2487

72. Rullo, O. J., Woo, J. M., Wu, H., Hoftman, A. D., Maranian, P., Brahn, B. A., McCurdy, D., Cantor, R. M., and Tsao, B. P. (2010) Association of IRF5 polymorphisms with activation of the interferon alpha pathway. Ann Rheum Dis 69, 611–617

73. Stone, R. C., Feng, D., Deng, J., Singh, S., Yang, L., Fitzgerald-Bocarsly, P., Eloranta, M. L., Ronnblom, L., and Barnes, B. J. (2012) Interferon regulatory factor 5 activation in monocytes of systemic lupus erythematosus patients is triggered by circulating autoantigens independent of type I interferons. Arthritis and rheumatism 64, 788–798

74. Yanai, H., Chen, H. M., Inuzuka, T., Kondo, S., Mak, T. W., Takaoka, A., Honda, K., and Taniguchi, T. (2007) Role of IFN regulatory factor 5 transcription factor in antiviral immunity and tumor suppression. Proceedings of the National Academy of Sciences of the United States of America 104, 3402–3407

75. Samuelson, L. C., and Metzger, J. M. (2006) Isolation and Freezing of Primary Mouse Embryonic Fibroblasts (MEF) For Feeder Plates. CSH Protoc 2006

76. Wen, F., Ellingson, S. M., Kyogoku, C., Peterson, E. J., and Gaffney, P. M. (2011) Exon 6 variants carried on systemic lupus erythematosus (SLE) risk haplotypes modulate IRF5 function. Autoimmunity 44, 82–89

77. Yanagiya, A., Svitkin, Y. V., Shibata, S., Mikami, S., Imataka, H., and Sonenberg, N. (2009) Requirement of RNA binding of mammalian eukaryotic translation initiation factor 4GI (eIF4GI) for efficient interaction of eIF4E with the mRNA cap. Molecular and cellular biology 29, 1661–1669

78. Wang, S., Wen, F., Wiley, G. B., Kinter, M. T., and Gaffney, P. M. (2013) An enhancer element harboring variants associated with systemic lupus erythematosus engages the TNFAIP3 promoter to influence A20 expression. PLoS Genet 9, e1003750

